# Postpartum resource scarcity reshapes maternal care, offspring prioritization, and the steroid-sensitive medial preoptic area

**DOI:** 10.64898/2026.11.09.765420

**Authors:** Dario Aspesi, Valeria Soriano, Sierra Rolle, Mohamed Y. Mahrous, Heather Brenhouse, Keerthi Krishnan, Debra A. Bangasser

**Affiliations:** Neuroscience Institute and the Center for Behavioral Neuroscience, Georgia State University, GA, 30303-5030, Atlanta, USA; Department of Biochemistry C Cellular and Molecular Biology, University of Tennessee at Knoxville, Knoxville, TN 37996, United States; Northeastern University, 125 Nightingale Hall, Boston, MA, 02115, USA

## Abstract

Parental care requires balancing offspring needs with environmental constraints, yet how resource scarcity alters the organization of maternal responses remains poorly understood. Here, we examined maternal behavior, offspring prioritization, and steroid receptor-associated extracellular matrix organization in postpartum rats exposed to limited bedding and nesting (LBN) or control conditions with ample nesting materials. LBN increased pup-directed and reduced self-directed behaviors but also increased behavioral entropy, a measure of behavioral fragmentation, indicating less predictable caregiving sequences. LBN dams preferentially retrieved male pups first, whereas control dams preferentially retrieved females first. LBN also increased the proportion of pup- investigation time directed toward males compared with controls. In offspring on postnatal day 10, corticosterone was elevated in female but not male LBN pups, a pattern that paralleled the maternal male bias. To identify associated neurobiological changes, we assessed steroid receptors and extracellular matrix organization in the medial preoptic area (mPOA), a key region for maternal behavior. LBN increased numbers of estrogen receptor α (ERα)- and androgen receptor (AR)- immunoreactive cells, reduced perineuronal net (PNN) abundance, and decreased receptor-positive cell enwrapment. Because PNNs constrain structural plasticity, these findings suggest that the postpartum environment alters the organization of steroid receptor-associated extracellular matrix, potentially modulating experience-dependent plasticity in neurosteroid-sensitive circuits. In sum, LBN increases the levels of maternal engagement while reshaping its temporal organization and allocation, accompanied by selective remodeling of steroid-sensitive preoptic architecture. Therefore, maternal adaptation to adversity may be better understood as changes in caregiving allocation strategies rather than a simple gain or loss of maternal care.

## Introduction

Postpartum resource scarcity is a challenge across species. In humans, food insecurity, unstable housing, limited social support, and related socioeconomic adversity are associated with poorer maternal mental health and child outcomes (Dunkel Schetter C Tanner, 2012; Kim et al., 2018). In other mammals, scarcity can limit access to food, shelter, or materials needed to protect and care for offspring. Motherhood is itself a highly plastic state, requiring rapid endocrine and neural adaptations that make pup cues motivationally salient and support nursing, nest attendance, licking and grooming, and retrieval (Barba-Müller et al., 2018; Bridges, 2015; Hillerer et al., 2014). These adaptations must sustain sufficient care while allowing mothers to meet their own energetic needs and respond to changing environmental demands. In rodents, the limited bedding and nesting (LBN) procedure isolates one component of adversity by restricting materials needed to construct and maintain a nest. Across studies, LBN produced either decreased or increased conventional measures of maternal care, including licking and grooming pups, nest attendance, and pup retrieval, depending on protocol implementation and species(Gallo et al., 2019; Ivy et al., 2008; Pardo et al., 2024; Rincón-Cortés C Grace, 2021; Walker et al., 2017) . However, studies examining behavioral sequences consistently report that LBN disrupts the temporal organization and predictability of maternal behavior (Demaestri et al., 2022; Eck et al., 2020; Gallo et al., 2019; Ivy et al., 2008; Molet et al., 2016; Pardo et al., 2024; Rincón-Cortés C Grace, 2021) . Distinguishing between the amount of care from its organization is critical: greater pup contact does not necessarily indicate stable or efficient caregiving.

Caregiving is also an allocation problem, requiring mothers to distribute finite time and energy between offspring care and self-maintenance and among individual offspring (Jensen Peña C Champagne, 2013; Macrì C Würbel, 2007; Veller et al., 2016). Dams must therefore decide which pups to prioritize, and these moment-to-moment decisions are guided by sensory cues from the pups themselves. Mothers integrate pup-derived olfactory and auditory signals during offspring search and retrieval, while pup movement, thermal state, and vocal output vary with physiological and environmental conditions and can shape maternal responses (Brunelli et al., 1994; Farrell C Alberts, 2002; Lenell et al., 2021; Moore, 1985; Nowlan et al., 2024; Stevenson et al., 2021). For example, mouse dams rearing all-male litters show greater nursing, nest building, or licking at specific postnatal ages than dams rearing mixed- or all-female litters (Alleva et al., 1989). Thus, individual offspring contribute multiple and potentially distinct sources of information to the caregiving environment and that dams can use this information when organizing maternal responses (Bauer, 1993; Gallo et al., 2019; Lapp et al., 2024; Numan et al., 2010). Maternal decisions therefore emerge from the integration of pup-related information with the broader environmental context. Resource scarcity could therefore alter not only whether and how consistently dams respond, but also how maternal responses are distributed among pups. Although LBN has previously been shown to impair pup retrieval, it remains unknown whether this reflects a generalized reduction in retrieval performance or a selective shift in offspring prioritization (Rincón-Cortés C Grace, 2021). Distinguishing these possibilities requires integrating measures of caregiving quantity and temporal structure with maternal choices among offspring.

LBN produces divergent long-term outcomes in male and female offspring, which could reflect sex-biased maternal allocation induced by low resources. In our previous work, LBN reduced impulsive choice and morphine self-administration in adult males, consistent with male-specific resilience to addiction-related phenotypes, whereas females did not show the same protection (Sanchez et al., 2021). In contrast, LBN disrupted cortical vasculature selectively in females, potentially compromising cortical function (Andrews et al., 2026). Maternal care can shape development of the offspring hypothalamic–pituitary–adrenal axis, and LBN effects on pup corticosterone vary across developmental stages and experimental protocols (Liu et al., 1997; Moussaoui et al., 2016; Walker et al., 2017). We therefore examined whether LBN produces sex- dependent changes in offspring corticosterone during the manipulation, providing a potential early endocrine correlate of divergent male and female trajectories.

The medial preoptic area (mPOA) is positioned to support flexibility in maternal behavior because it integrates pup-related sensory information with reproductive and energetic state and organizes multiple components of parental care (Jamieson C Kohl, 2026; Numan C Insel, 2006). Estrogen receptor alpha (ERα)-expressing mPOA neurons are recruited during pup retrieval and are required for its normal expression (Wei et al., 2018). In lactating rats, natural variation in licking and grooming is associated with ERα expression in the mPOA, maternal interactions activate ERα- positive neurons, and differences in maternal care covary with the dendritic complexity of mPOA neurons (Champagne et al., 2003; Lonstein et al., 2000; Parent et al., 2017). Intriguingly, androgen receptor (AR)-expressing neurons are also abundant in the mPOA; in male mice, AR- and ERα- expressing cells share several neuropeptide identities, but the organization of AR-positive populations in the maternal brain is poorly understood (Tsuneoka et al., 2017). Resource scarcity may therefore engage steroid-sensitive mPOA populations, although evidence for postpartum remodeling of these populations is limited.

Perineuronal nets (PNNs) also provide a candidate structural substrate for the behavioral effects of LBN on dams. These specialized extracellular matrix structures stabilize mature circuits and can constrain experience-dependent synaptic change; disrupting them can reopen plasticity in adult circuits (Carstens et al., 2021; Grieb et al., 2021; Hylin et al., 2013; Lau et al., 2020; Lupori et al., 2023; Pizzorusso et al., 2002). Changes in PNN abundance may influence not only whether maternal circuits are plastic, but also how flexibly mothers can adjust behavioral strategies when offspring demands or environmental conditions change. In the rat mPOA, PNNs assemble during pregnancy, enwrap ERα-positive neurons, and decline during lactation (Uriarte et al., 2020), indicating that extracellular matrix remodeling accompanies the transition to motherhood. Whether resource scarcity further alters PNN abundance or their distribution across ERα- and AR-expressing cells is unknown.

Here, we tested whether LBN alters (1) the amount and temporal predictability of maternal care, (2) retrieval efficiency and the allocation of responses toward male and female pups, (3) ERα and AR immunoreactivity and receptor-associated PNN organization in the mPOA, and (4) offspring plasma corticosterone levels. We hypothesized that LBN would increase pup-directed care while reducing its temporal predictability and delaying pup retrieval. We further predicted that LBN would alter the allocation of maternal investigation between male and female pups. In parallel, we examined whether these behavioral changes were accompanied by altered steroid receptor immunoreactivity and receptor-associated PNN organization in the mPOA.

### Material and Methods Subjects

Adult female Long–Evans rats (approximately 2 months old) were purchased from Charles River Laboratories (Wilmington, MA, or Kingston, NY, USA). Following at least 1 week of acclimation, females were housed with male breeders for 2 weeks and subsequently separated from the males for the remainder of pregnancy. All dams were primiparous. The day of birth was designated postnatal day (PND) 0. On PND2, litters were culled to 10 pups, with five males and five females whenever possible.

On PND2, dams and litters were randomly assigned to either control (CTRL) or limited bedding and nesting (LBN) housing in standard shoebox cages fitted with filter tops (Fig. 1A). Control cages contained standard nesting material, direct access to Bed-o’Cobs bedding, and an enrichment tube. LBN cages contained one paper towel for nesting, no enrichment, and a metal grate that prevented direct access to the bedding below (Andrews et al., 2026; Eck et al., 2020; Ordoñes Sanchez, 2021). Food and water were available *ad libitum* throughout the experiment, except for LBN dams, which received 50g of food on PND2 that was not refilled to reduce the use of food as nesting material. The amount of food provided was based on previous observations and was enough for free feeding through the duration of the LBN protocol. Animals received LabDiet 5001 chow, whereas breeding pairs received LabDiet 5015 chow (LabDiet, St. Louis, MO, USA). All animals were housed in the Central Animal Facility at Georgia State University under a reversed 12:12-h light–dark cycle, with lights off at 10:00 a.m.

**Figure 1.**
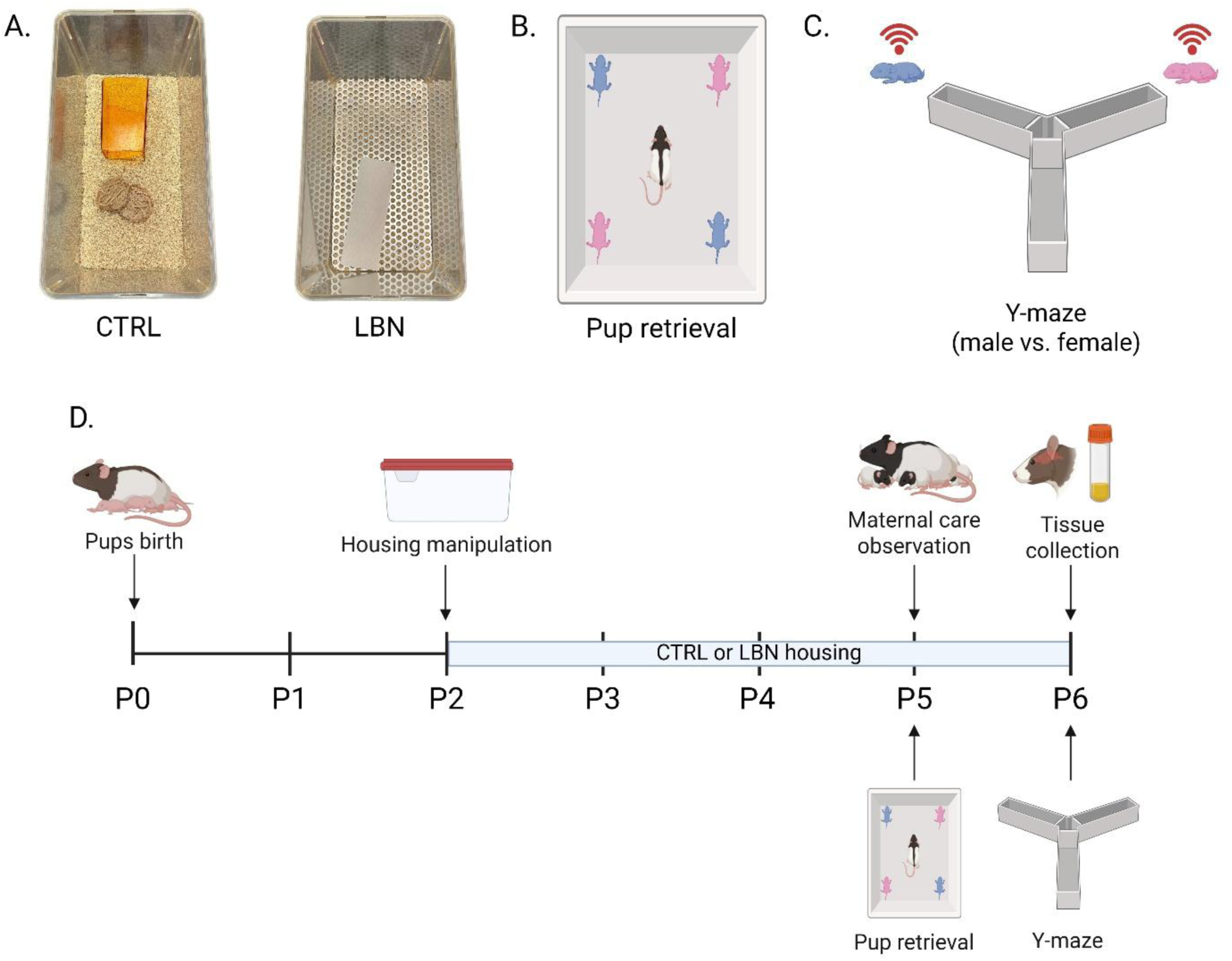
Experimental design and behavioral paradigms. (A) On postnatal day (PND) 2, dams and litters were randomly assigned to control (CTRL) or limited bedding and nesting (LBN) housing. CTRL cages contained standard bedding, nesting material, and an enrichment tube, whereas LBN cages contained a perforated metal platform preventing direct access to bedding, one paper towel for nesting, and no enrichment. Maternal behavior was recorded twice daily from PND2–6, with PND5 sessions used for analysis. (B) Pup retrieval was assessed on PND5 by placing four littermates, two males and two females, in the corners of the home cage and recording retrieval latency and order. (C) On PND6, dams completed Y-maze choice tests comparing responses to one of their own male and female pups. Stimulus pups were placed beneath perforated enclosures that permitted auditory and olfactory access while preventing direct contact and retrieval. (D) Schematic representation of the experimental design. Dam brains were collected after PND6 testing; pup plasma was obtained from independent PND6 and PND10 cohorts for corticosterone analysis. Blue and pink denote male and female pups, respectively.

Unless otherwise indicated, dams and litters remained in their assigned housing conditions until PND6, when the animals were euthanized and tissues were collected. Maternal behavior was analyzed in 29 dams (CTRL, n = 14; LBN, n = 15). Different subsets of dams contributed to the subsequent behavioral and neuroanatomical experiments (Fig. 1B-D). Pup retrieval included 13 CTRL and 13 LBN dams. A separate cohort used for the male-versus-female Y-maze experiment included 11 CTRL and 10 LBN dams. ERα/PNN analyses were performed in 6 CTRL and 6 LBN dams, whereas AR/PNN analyses included 5 CTRL and 6 LBN dams.

Plasma corticosterone was assessed in offspring using an independent sample assembled from litters generated across multiple experiments in the laboratory, including but not limited to the maternal-behavior cohorts described above. At PND6, samples were obtained from 17 CTRL litters and 18 LBN litters; at PND10, samples were obtained from 16 CTRL litters and 18 LBN litters. No litter contributed more than one pup of one sex to either corticosterone analysis, and each analyzed pup therefore represented an independent litter. Detailed sex-specific sample sizes are reported in the Corticosterone ELISA measurement section.

All procedures were approved by the Georgia State University Institutional Animal Care and Use Committee and were conducted in accordance with the National Institutes of Health *Guide for the Care and Use of Laboratory Animals*.

### Maternal Behavior

The LBN protocol has been shown to disrupt and fragment maternal behavior (Ivy et al., 2008; Walker et al., 2017). Maternal behavior was video-recorded twice daily in 1-h sessions. The first session began 4 h before lights off, during the light phase, and the second began 5 h before lights on, during the dark phase (timed to minimize entry into the room by staff). Recordings were subsequently scored offline. Although recordings were collected throughout the LBN manipulation, the present analyses focused on PND5 and both daily sessions (light cycle and dark cycle) as this is where we and others have seen the largest differences in care between groups (Gallo et al., 2019; Ivy et al., 2008; Molet et al., 2016; Moussaoui et al., 2016; Pardo et al., 2024). Both daily sessions analyzed separately and averaged within each dam. This time point was selected based on previous studies identifying PND5 as a sensitive period for detecting experimentally induced alterations in maternal behavior (Eck et al., 2020; Grieco et al., 2026; Lauraine et al., 2024; Martín-Sánchez et al., 2015; Pardo et al., 2024; Sarkisova C Gabova, 2018). Maternal behavior was recorded from all dams, but different subsets of animals were exposed to the other behavioral paradigms. Pups were not individually marked or tracked during home-cage recordings; therefore, maternal care was quantified at the litter level rather than for individual pups.

Scored behaviors included nursing, licking and grooming pups, nest building, self-grooming, returning displaced pups to the nest, locomotor activity, eating, drinking, and resting away from the nest. For composite analyses, nursing, licking and grooming pups, and returning pups to the nest were classified as active pup-directed behaviors. Self-grooming, eating, drinking, and resting away from the nest were classified as self-directed behaviors. Nest building and locomotor activity were analyzed separately. The frequency, total duration, and mean bout duration of each behavior were recorded, where applicable. Videos were scored using BORIS behavioral observation software (v.7.13.9) (Friard C Gamba, 2016).

Once all the scores were obtained, an entropy statistic was calculated based on a mathematical computational code provided by Vegetabile et al. (Vegetabile et al., 2019). The entropy statistic uses the conditional probability of each type of behavior following other types of behavior in a sequence, to determine whether the behavior of the dams during LBN treatment is unpredictable. Higher entropy values indicate a less predictable behavioral sequence. This entropy score has been used by a variety of research groups to validate the treatment and effectiveness of the LBN model (L. K. Davis et al., 2020; Demaestri et al., 2022; Jones et al., 2025; Walker et al., 2017).

To visualize differences in behavioral organization, transitions among behaviors were also modeled as first-order, discrete-time Markov chains using a lag of one behavioral event. Transition- network analyses were performed as previously described (Saqr et al., 2025; Stevenson et al., 2021) using the *behavseq* package, and sequence visualizations were generated using the *TraMineR* package (version 2.2). (*GitHub - Jalapic/Behavseq: Sequential Methods for the Behavioral Sciences · GitHub*, n.d.). Only transitions with conditional probabilities ≥ 0.20 are shown in the representative networks.

### Pup Retrieval

Pup retrieval, a behavioral procedure used to assess maternal motivation and caregiving, was assessed on PND5 at the beginning of the dark phase. Dams and litters were transferred to the behavioral testing room and allowed to habituate for at least 30 min. At the beginning of the test, all pups were removed from the home cage and maintained on a heating pad to minimize heat loss.

Following a 5-min separation period, four pups (two males and two females) were returned to the home cage, with one pup placed in each corner (Fig. 1B). Male and female pups were systematically counterbalanced or randomized across cage corners. Maternal behavior was video-recorded, and the latency to retrieve each pup to the nest was scored. The order and sex of the retrieved pups were also recorded. Based on pilot experiments indicating that dams typically retrieved pups rapidly, the test was limited to 5 min. Dams that did not complete a given retrieval within the 300-s observation period were right-censored at 300 s. BORIS software was used to score retrieval latencies and retrieval order, as well as the duration of licking and grooming pups and locomotor activity during the test.

### Y-maze pup-cue preference

Maternal responses to multimodal cues from pups belonging to the dam’s own litter were assessed on PND6 using a Y-maze apparatus, adapting the protocol previously described in (Rekapalli et al., 2024). The rat Y-maze (San Diego Instruments, San Diego, CA, USA) consisted of three arms, each measuring 55.9 cm long and 17.8 cm wide, with 10.2-cm-high walls. The starting compartment was 25.4 cm long. All testing was conducted under red-light illumination. On the day of testing, dams were transferred to the behavioral testing room and allowed to habituate for at least 30 min.

To test for sex preference, a male pup and a female pup from the dam’s litter were placed on a weigh boat beneath a perforated 3D-printed enclosure in the two goal arms. The enclosure allowed the dam access to auditory and olfactory cues while preventing direct contact with or retrieval of the pup. The locations of the male and female pups were counterbalanced across dams. The dam was placed in the third, starting arm of the Y-maze, and its behavior was recorded for 5 min. The Y-maze and 3D-printed enclosures were cleaned with 70% ethanol in water and allowed to dry completely between dams.

Videos were scored using BORIS software by observers blind to housing conditions. Outcome measures included the duration of time spent in the central area of the Y-maze, the number of entries into each goal arm (entry occurred when all four paws crossed into a goal arm), latency to enter each arm, total time spent in each arm, and duration of active pup investigation. Active investigation was defined as sniffing through or biting the perforations of the enclosure. An investigation preference score was calculated by dividing the time spent investigating the male pup by the total time spent investigating both the male and female pups. In both experiments, a score of 0.5 indicated no preference.

### Tissue collection

On PND6, following completion of the Y-maze test, dam brain tissue was collected for analyses. Pup plasma was collected on PND6 or, for the second cohort, on PND10. All brain tissue and plasma collections were performed between 11:00 a.m. and 1:00 p.m., corresponding to 1–3 h after lights off at 10:00 a.m. Dams were deeply anesthetized with an overdose of sodium pentobarbital and transcardially perfused. The thoracic cavity was opened through abdominal and transdiaphragmatic incisions, the right atrium was incised, and a perfusion needle was inserted into the left ventricle. Animals were perfused with ice-cold phosphate-buffered saline (PBS) to remove circulating blood, followed by 4% paraformaldehyde (PFA) in 0.1 M PBS (pH 7.4). Brains were removed and postfixed in 4% PFA for 24 h at 4°C, followed by cryoprotection for 24–48 h in PBS containing 30% sucrose. Brains were subsequently flash-frozen in isopentane precooled to approximately −30 to −40°C over dry ice and stored at −80°C. The day before sectioning, brains were transferred to a −20°C freezer. Coronal sections were collected at a thickness of 40 µm using a Leica CM3050 S cryostat. Free-floating sections were stored at −20°C in multiwell plates containing antifreeze solution (30% glycerol, 30% ethylene glycol, 30% deionized water, and 10% phosphate solution containing 0.0243 M NaH₂PO₄ and 0.0175 M NaOH).

On PND6 or PND10, pups were rapidly decapitated, and trunk blood was collected into 500- μl EDTA-coated BD Microtainer tubes (Cat. No. 22-253-270; Fisher Scientific, USA). Samples were centrifuged at 3,000 × *g* for 15 min at 4°C; plasma was transferred to 0.5-ml tubes and stored at −80°C. Although two male and two female pups were collected from each litter, only one pup per litter was randomly selected for corticosterone analysis, while balancing pup sex across housing conditions. Samples were selected to balance sex and housing condition as closely as possible, and no litter contributed more than one analyzed pup.

### Double-label immunofluorescence: ERα/PNN and AR/PNN

To determine whether LBN affected steroid receptor expression, PNN abundance, and the spatial association between steroid receptor-positive cells and PNNs, separate series of mPOA sections were processed for ERα/WFA or AR/WFA double-label immunofluorescence. WFA is an N- acetylgalactosamine-binding lectin commonly used to label PNN-associated extracellular matrix structures (Härtig et al., 2022; Lupori et al., 2023). Sections spanning approximately −0.20 to −1.20 mm relative to bregma were identified using a rat brain atlas (Paxinos C Watson, 2013). Because detectable WFA-positive PNNs were restricted to four or five sections per animal, analyses were performed on these sections. The same rostrocaudal levels were used to quantify ERα and AR expression, although receptor immunoreactivity extended across a broader portion of the mPOA.

Free-floating sections were washed five times in PBS and blocked for 1 h in PBS (pH 7.4) containing 0.2% Triton X-100, 5% normal donkey serum, and 1% bovine serum albumin. Separate section series were then incubated for 24 h at 4°C in one of two primary-labeling cocktails: rabbit anti-ERα antibody (1:500; Cat. No. 06-935, Millipore Sigma; (Aspesi et al., 2026; Zou et al., 2025)) and fluorescein-labeled *Wisteria fforibunda* agglutinin (WFA;1:200; Cat. No. FL-1351-2; Vector Labs, Burlingame, CA) or rabbit anti-AR antibody (1:250; Cat. No. ab133273, Abcam; (Aspesi et al., 2026)) and fluorescein-conjugated WFA at the same concentration. The following day, sections were washed in PBS and incubated for 1 h with an Alexa Fluor 594-conjugated donkey anti-rabbit IgG secondary antibody (1:400; Cat. No. 711-585-152, Jackson ImmunoResearch Laboratories, USA) diluted in blocking solution. No secondary detection was required for directly conjugated WFA. Sections were subsequently washed in PBS, mounted onto microscope slides, and coverslipped using VECTASHIELD HardSet Antifade Mounting Medium with DAPI (Vector Labs, Burlingame, CA).

Negative controls included sections processed with omission or replacement of the primary antibody with an equivalent concentration of normal serum, as well as sections processed without the secondary antibody.

### Quantitative analysis

Images were acquired using a Leica DM4 B microscope with a 10× objective under identical acquisition settings across experimental groups. Images were analyzed using ImageJ/Fiji software (version 1.54t; National Institutes of Health, Bethesda, MD, USA) by an analyst blind to treatments. WFA-positive PNNs and associations between PNNs and steroid receptor-positive cells were quantified manually using the “Cell Counter” plugin.

Because of the high density of ERα- and AR-positive cells in the mPOA, receptor-positive cells were quantified using an intensity-thresholding procedure. Threshold limits were adjusted individually for each section to include clearly identifiable receptor-positive labeling while excluding background fluorescence. All analyses were conducted within a predetermined mPOA region of interest measuring approximately 2.07 mm². Counts obtained from the analyzed sections were averaged to produce one value per animal, which served as the experimental unit.

To determine whether LBN affected the cellular distribution of PNNs in addition to their overall abundance, two complementary measures of receptor–PNN association were calculated. The percentage of receptor-positive cells surrounded by PNNs was calculated as: (receptor-positive cells surrounded by a PNN / total receptor-positive cells) × 100. The percentage of PNNs surrounding receptor-positive cells was calculated as: (PNNs surrounding a receptor-positive cell / total PNNs) × 100. A receptor-positive cell was considered surrounded by a PNN when a receptor-positive cell was fully enwrapped by a PNN. Cells labeled for steroid receptors in close proximity to PNNs were not counted in this analysis. Figure 2 and Supplementary video 1 illustrate representative examples of this classification.

**Figure 2.**
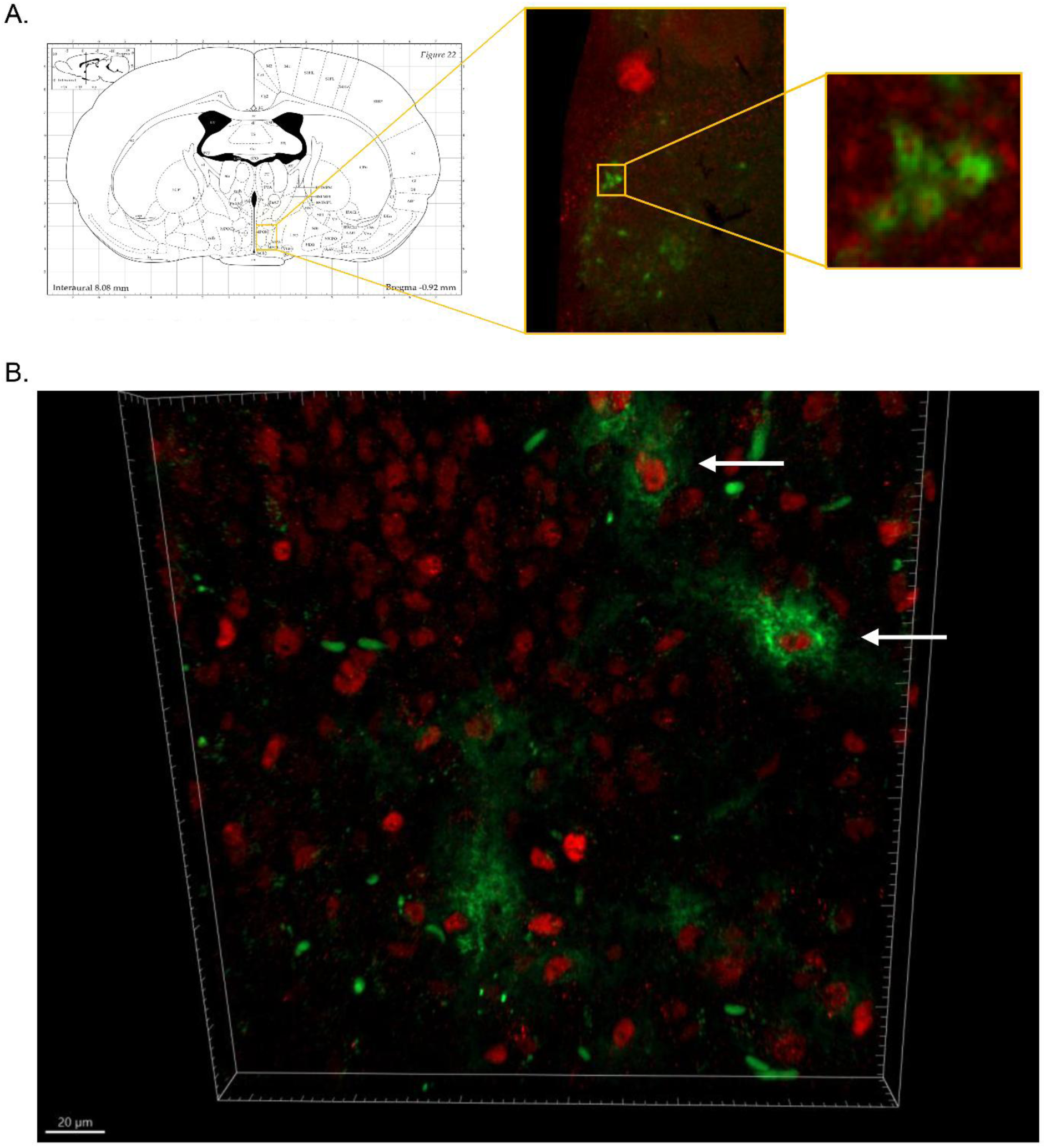
Representative visualization of ERα-positive cells and perineuronal nets in the mPOA. Coronal atlas diagram indicating the medial preoptic area (mPOA) sampling region (yellow box), accompanied by a representative dual-label fluorescence image and an enlarged view illustrating the spatial association between an ERα-positive cell and a Wisteria floribunda agglutinin (WFA)- labeled perineuronal net (PNN). (B) Three-dimensional reconstruction of a fluorescence z-stack showing ERα immunoreactivity (red) and WFA-labeled PNNs (green). White arrows indicate ERα- positive cells enwrapped by PNNs. Scale bar, 20 μm.

### ELISA Corticosterone analysis

Plasma corticosterone concentrations were measured in duplicate using a commercially available ELISA kit according to the manufacturer’s instructions (Corticosterone ELISA Kit, catalog no. ADI-900-097; Enzo Life Sciences, Farmingdale, NY, USA). Technical duplicates were averaged to obtain a single corticosterone value for each pup. Samples were distributed across the assay to balance housing condition and sex as closely as possible. Each analyzed pup originated from a different litter and therefore represented an independent biological replicate. Absorbance was measured at 405nm, using a Bio-Rad iMark spectrophotometer plate reader. The mean intra-assay coefficient of variation was 9.1%. Consequently, each analyzed value represented an independent litter.

### Statistical Analysis

Statistical analyses were performed using GraphPad Prism version 11 (GraphPad Software, Boston, MA, USA) and R version 4.4.1. All statistical tests were two-tailed. Normality and homogeneity of variance were evaluated using Shapiro–Wilk and Brown–Forsythe tests, respectively, together with visual inspection of Q–Q plots.

Comparisons between CTRL and LBN dams for behavioral and morphological outcomes were conducted using independent-samples t-tests when parametric assumptions were met and Mann–Whitney U tests otherwise. In the Y-maze experiment, the proportion of total pup-investigation time directed toward the male pup and the proportion of total goal-arm time spent in the male- associated arm were compared between housing conditions using independent-samples t-tests.

The time required to retrieve all four pups was analyzed using Kaplan–Meier time-to-event analysis. Retrieval of the fourth pup was defined as completion of the retrieval sequence, and dams that did not retrieve all four pups during the 300-s testing period were right-censored at 300 s. Completion-time distributions for CTRL and LBN dams were compared using the log-rank Mantel– Cox test. The association between housing condition and the sex of the first pup retrieved was evaluated using Fisher’s exact test. Later retrieval positions were not analyzed as independent pup- sex choices because each retrieval altered the sex composition of the pups remaining in the cage.

Plasma corticosterone concentrations were analyzed separately at PND6 and PND10 using ordinary two-way ANOVAs, with housing condition and pup sex as between-subjects factors. Because every analyzed pup originated from a different litter, each observation represented an independent biological replicate. Significant interactions were followed by Šídák-corrected multiple- comparison tests.

Associations between behavioral and morphological measures were assessed using Pearson correlations. Simple linear regression was used to visualize relationships between variables, and differences in regression slopes between housing conditions were assessed using models containing the interaction between housing condition and the continuous predictor. Corrections for multiple correlations or slope comparisons were applied as specified for each family of analyses.

Effect sizes for t tests and ANOVAs are reported as eta squared (η²), with values of 0.01, 0.06, and 0.14 interpreted as small, medium, and large effects, respectively (Cohen, 2013). Effect sizes for Mann–Whitney U tests are reported as rank-biserial correlations (r_rb_), with absolute values of 0.10, 0.30, and 0.50 interpreted as small, medium, and large effects, respectively (Cohen, 2013). For time- to-event analyses, median completion times and the proportions of animals completing the retrieval sequence are reported. Data are presented as mean ± SEM unless otherwise indicated. Individual data points represent animals or litter-level means, as specified in the figure legends. For experiments involving dams, n indicates the number of dams. For pup analyses, n indicates the number of independent litters. Statistical significance is denoted in the figures as follows: *p < .05, **p < .01, ***p <.001, and ****p < .0001.

## Results

### LBN increases maternal care but reduces the predictability of behavioral sequences

On average across light and dark phase on PND5, LBN dams spent more time engaged in pup-directed behaviors than CTRL dams (t(27) = 3.36, p < .01, η² = .29; Fig. 3A) and less time engaged in self-directed behaviors (t(27) = 3.01, p < .01, η² = .25; Fig. 3B). Analysis of individual behaviors showed that LBN dams spent more time nursing (t(27) = 2.59, p < .05, η² = .20; Fig. 3D) and carrying pups (U = 26, p < .001, η² = .32; Fig. 3E), while spending less time away from the nest (t(27) = 6.97, p < .0001, η² = .64; Fig. 3G). No significant group differences were detected for the other behaviors included in the composite measures (all p > .05).

**Figure 3.**
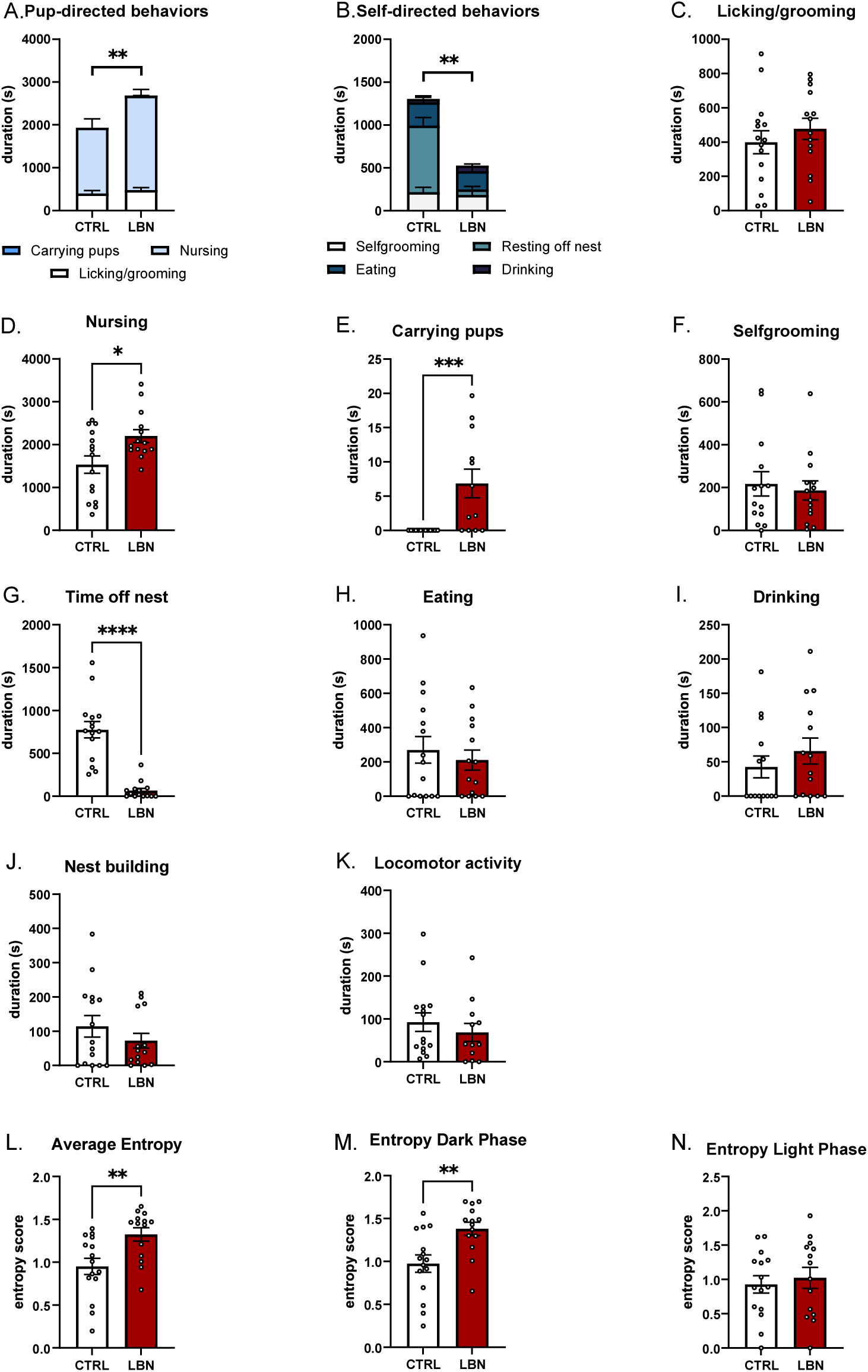
LBN increases pup-directed maternal care and reduces the predictability of behavioral sequences. (A) Total duration of pup-directed behaviors, comprising licking/grooming, pup carrying, and nursing. (B) Total duration of self-directed behaviors, comprising self-grooming, eating, drinking, and resting off the nest. (C–K) Duration of individual behaviors: licking/grooming (C), nursing (D), pup carrying (E), self-grooming (F), time off the nest (G), eating (H), drinking (I), nest building (J), and locomotor activity (K). (L–N) Behavioral entropy averaged across recording phases (L) and during the dark (M) and light (N) phases. Higher entropy scores indicate less predictable behavioral sequences. Bars represent mean ± SEM, and individual points represent dams. CTRL, control housing; LBN, limited bedding and nesting. *p < .05, **p < .01, ***p < .001, ****p < .0001.

Despite this increase in pup-directed care, LBN dams exhibited higher behavioral entropy than CTRL dams (t(27) = 3.03, p < .01, η² = .25; Fig. 3L), indicating that their behavioral sequences were less predictable. When the light and dark phases were examined separately, the group difference was significant during the dark phase (t(27) = 3.16, p < .01, η² = .27; Fig. 3M) but not during the light phase (t(27) = 0.48, p = .63, η² = .008; Fig. 3M).

Transition-probability networks provided a complementary visualization of these differences in behavioral organization (Fig. 4). Transitions in LBN dams were distributed across a broader range of behavioral states, whereas the CTRL network was dominated by fewer, more recurrent transitions. Thus, LBN increased the overall amount of pup-directed care while reducing the stability and predictability of its temporal organization.

**Figure 4.**
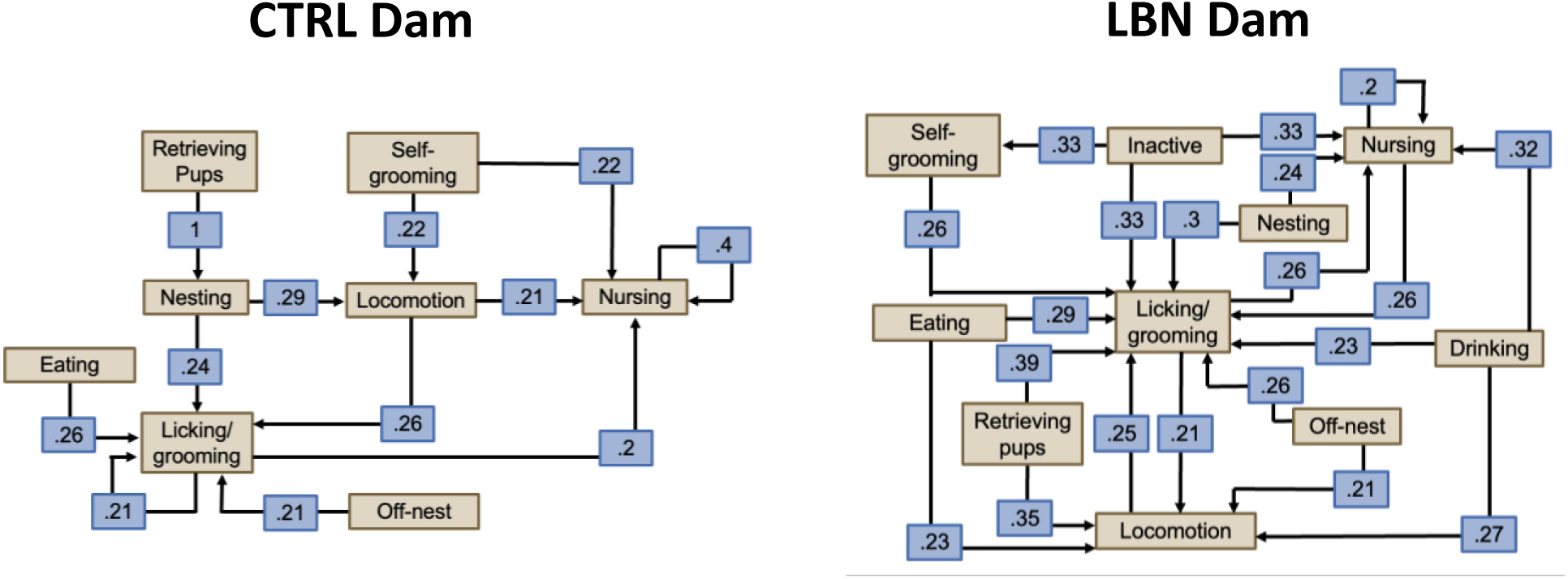
Representative transition networks illustrate altered maternal behavioral sequencing following LBN. Representative lag-1 transition-probability networks generated from PND5 maternal behavior sequences for a CTRL dam (left) and an LBN dam (right). Beige boxes represent observed behavioral states, directed arrows indicate transitions from one behavior to the next, and blue boxes report the conditional probability of each transition. Compared with the CTRL example, the LBN network contains a broader range of behavioral states and interconnected transition pathways, visually illustrating the greater fragmentation of maternal behavior under LBN conditions. Networks are representative of the behavioral sequences included in the entropy analysis; group analyses included 14 CTRL and 15 LBN dams. CTRL, control housing; LBN, limited bedding and nesting.

### LBN delays completion of pup retrieval and shifts maternal pup-cue responsivity toward male offspring

All CTRL dams completed retrieval of the four pups within the 300-s testing period (13/13), compared with 7 of 13 LBN dams. The six LBN dams that did not complete the retrieval sequence were right-censored at 300 s. Kaplan–Meier analysis showed that retrieval-completion times differed between housing conditions (log-rank Mantel–Cox test, χ²(1) = 12.39, p < 0.001; Fig. 5A). Median completion time was 178.85 s in CTRL dams and 253.25 s in LBN dams. Consistent with the time-to- event analysis, a complementary Mann–Whitney U sensitivity analysis, in which censored observations were retained at the 300-s ceiling, showed longer fourth-pup latencies in LBN than CTRL dams (U = 23, p = .0016, r_rb_ = 0.73). Together, these analyses indicate that LBN delayed completion of the retrieval sequence.

**Figure 5.**
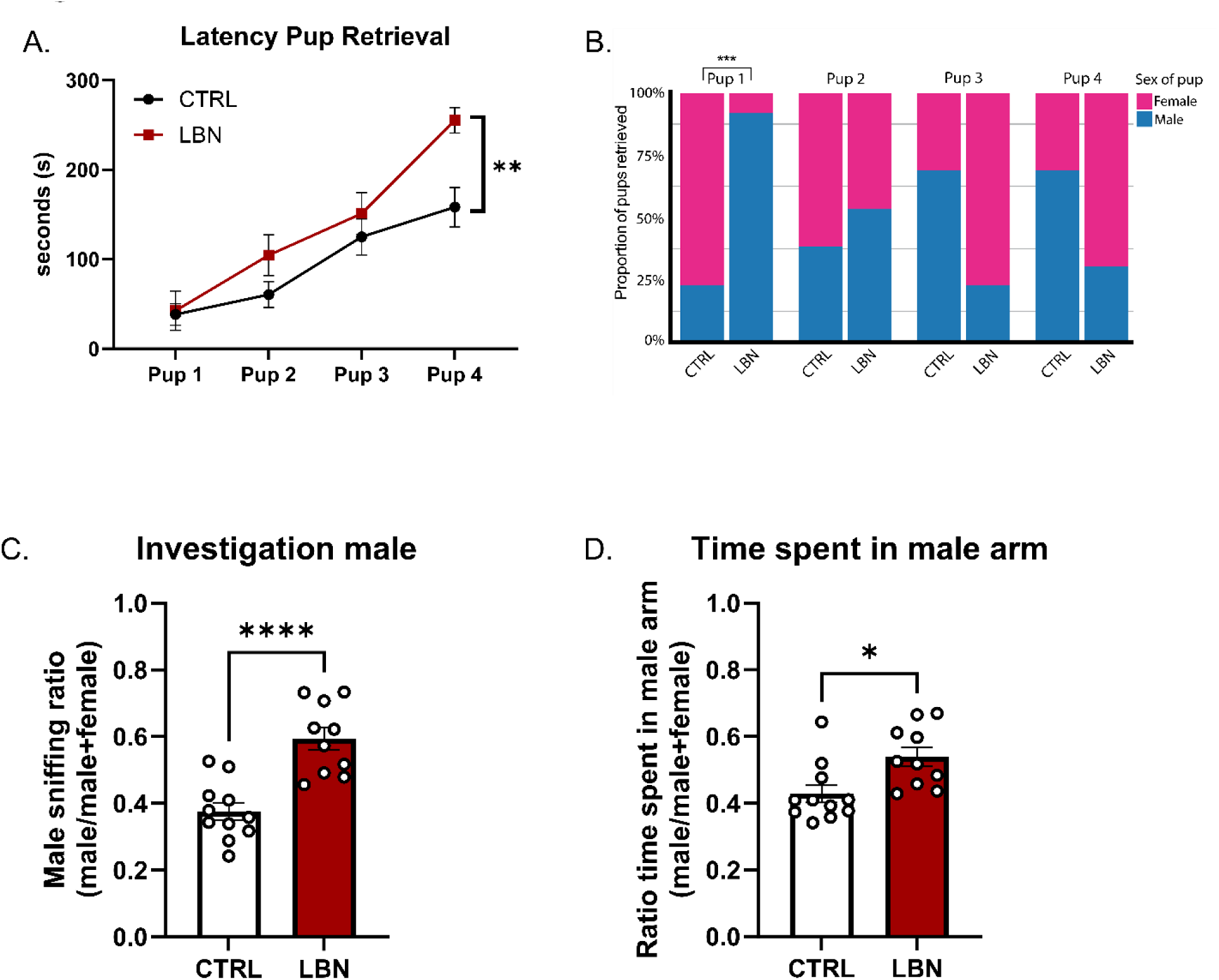
LBN delays completion of pup retrieval and shifts maternal responses toward male offspring. (A) Cumulative latency to retrieve four pups (two males and two females) on postnatal day (PND) 5. LBN dams took longer to retrieve the fourth pup. (B) Proportions of male (blue) and female (pink) pups retrieved at each position. Later retrieval positions are constrained by the sex of previously retrieved pups. (C, D) In a separate cohort of dams, one male pup and one female pup were presented simultaneously in different arms of a Y-maze, and we quantified the proportion of total pup-investigation time directed toward the male (C) and the proportion of total goal-arm time spent in the male-associated arm (D). A score of 0.5 indicates equal allocation in panels C and D. Data represent mean ± SEM, except in B, where bars show proportions; individual points in C and D represent dams. n = 13 dams per group (A and B); n = 11 control and 10 LBN dams (C and D). Asterisks indicate between-group differences: *p < .05, **p < .01, ***p < .001, ****p < .0001. CTRL, control housing; LBN, limited bedding and nesting.

The sex of the first pup retrieved also differed between housing conditions (Fisher’s exact test, p = .001; Fig. 5B). CTRL dams retrieved a female pup first in 10 of 13 trials (76.9%), whereas LBN dams retrieved a male pup first in 12 of 13 trials (92.3%). Although the sex distributions also differed at later retrieval positions, these comparisons were not independent because each dam was presented with two pups of each sex and every retrieval constrained the sex composition of the remaining pups. Therefore, the first retrieval provides the clearest evidence that LBN shifted initial pup prioritization toward male offspring.

In the Y-maze experiment, dams were simultaneously presented on of their own male pup and female pups. The male-pup investigation score was higher in LBN dams than in CTRL dams (t(19) = 5.15, p < .001, η² = .58; Fig. 5E). LBN dams also allocated a greater proportion of their total goal- arm time to the male-associated arm than CTRL dams (t(19) = 2.84, p < .05, η² = .30; Fig. 5F). No group differences were detected for the remaining behavioral outcomes (all p > 0.05). Together, these findings indicate that LBN shifted maternal responsivity toward male-related pup cues.

### LBN differentially reorganizes PNN associations with steroid receptor–expressing cells in the mPOA

#### LBN increases ERα-positive cells while reducing their PNN enwrapment in the mPOA

In the mPOA, LBN dams showed a significant increase in the number of ERα-positive cells (t(10) = 2.75, p < .05, η² = .43; Fig. 6A), but fewer PNNs (t(10) = 2.38, p < .05, η² = .36; Fig. 6B) compared with CTRL dams. When evaluating the percentage of ERα-positive cells surrounded by PNNs, LBN dams showed a significant decrease (t(10) = 3.82, p < .01, η² = .59; Fig. 6C), consistent with the concurrent increase in steroid receptor-expressing cells and decrease in PNN abundance. However, the percentage of PNNs surrounding ERα-positive cells did not differ between groups (t(10) = 1.34, p = .21, η² = .15; Fig. 6D).

**Figure 6.**
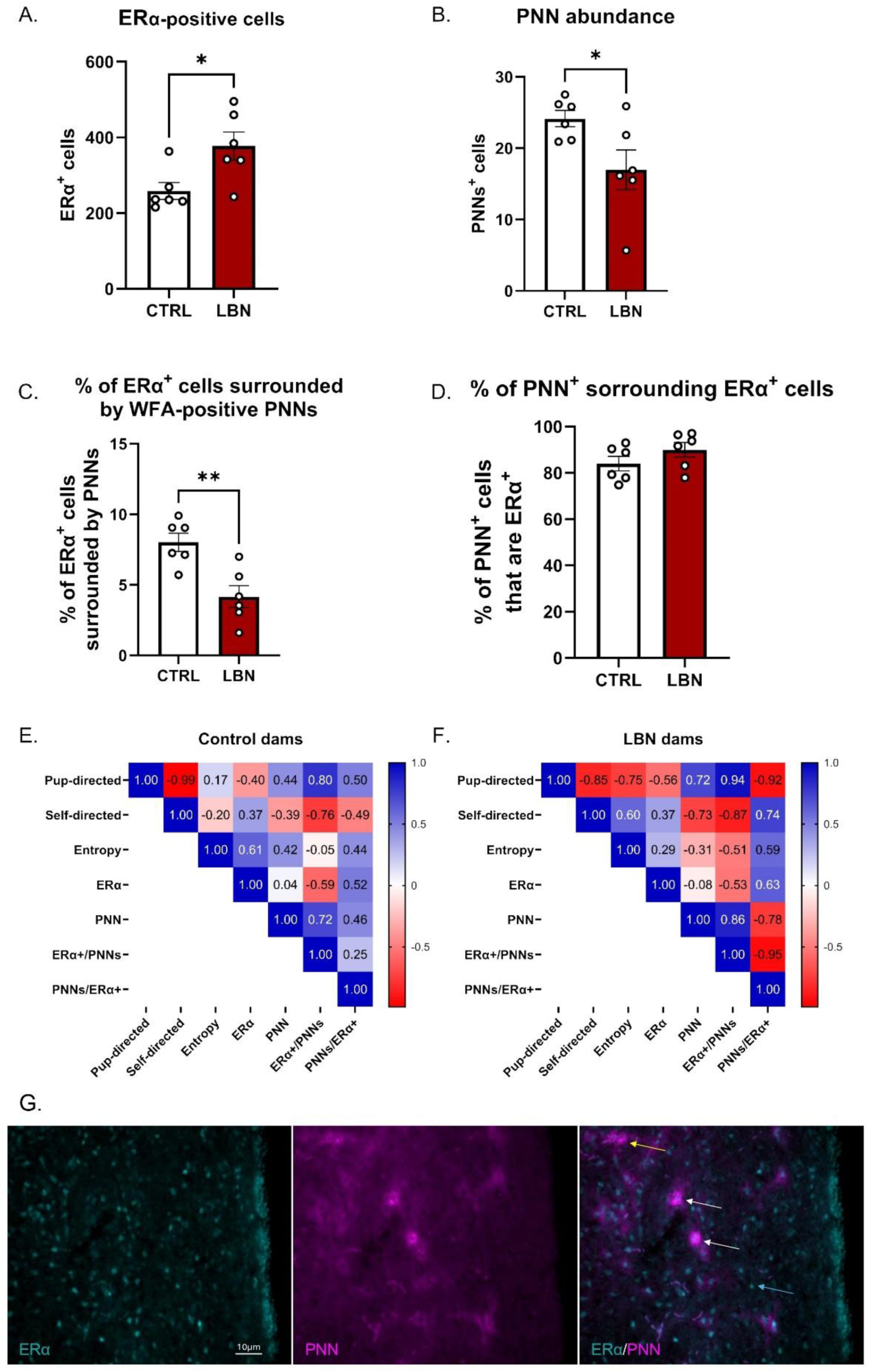
LBN increases ERα-positive cells while reducing PNN abundance and ERα-cell enwrapment in the mPOA. (A) Number of estrogen receptor α-positive (ERα+) cells in the medial preoptic area (mPOA). (B) Number of Wisteria floribunda agglutinin-labeled perineuronal nets (PNNs). (C) Percentage of ERα+ cells surrounded by PNNs. (D) Percentage of PNNs surrounding ERα+ cells. LBN increased ERα+ cell number while reducing overall PNN abundance and the proportion of ERα+ cells enwrapped by PNNs. The proportion of the remaining PNNs associated with ERα+ cells did not differ between groups. (E, F). Exploratory Pearson correlation matrices showing associations among pup-directed behavior, self-directed behavior, behavioral entropy, ERα+ cell number, PNN abundance, the percentage of ERα+ cells surrounded by PNNs, and the percentage of PNNs surrounding ERα+ cells in control (E) and LBN dams (F). (G) Representative fluorescence micrographs of ERα immunoreactivity (cyan; left), WFA-labeled PNNs (magenta; center), and merged ERα/PNN labeling (right) in the mPOA. In the merged image, white arrows indicate ERα-positive cells enwrapped by WFA-positive PNNs, light-blue arrows indicate ERα-positive cells without detectable PNN enwrapment, and yellow arrows indicate WFA-positive PNNs not enwrapping ERα-positive cells. Values represent Pearson correlation coefficients (r); blue indicates positive associations and red indicates negative associations. Bars represent mean ± SEM, and individual points represent dams. n = 6 dams per group. Group comparisons in A–D were performed using independent-samples t- tests. *p < .05; **p < .01. CTRL, control housing; LBN, limited bedding and nesting.

Correlation analyses were conducted separately in CTRL and LBN dams to explore associations among maternal behavior, ERα expression, PNN abundance, and ERα–PNN organization in the mPOA (Fig. 6E, F). In CTRL dams, pup-directed behavior was positively associated with the percentage of ERα-positive cells surrounded by PNNs (r = .80, p = .056, n = 6), whereas self- directed behavior showed a negative association with the same measure (r = −.76, p = .079, n = 6). The remaining behavioral–morphological associations were weak to moderate.

A different pattern emerged in LBN dams. Pup-directed behavior was strongly positively associated with the percentage of ERα-positive cells surrounded by PNNs (r = .94) and strongly negatively associated with the percentage of PNNs surrounding ERα-positive cells (r = −.92). Conversely, self-directed behavior was negatively associated with the percentage of ERα-positive cells surrounded by PNNs (r = −.87) and positively associated with the percentage of PNNs surrounding ERα-positive cells (r = .74). Pup-directed and self-directed behaviors also showed opposing associations with overall PNN abundance (r = .72 and r = −.73, respectively). In contrast, behavioral entropy showed only weak-to-moderate associations with the morphological measures in LBN dams (r ≤ 0.59). Within the LBN group, PNN abundance was positively associated with the percentage of ERα-positive cells surrounded by PNNs (r = .86) and negatively associated with the percentage of PNNs surrounding ERα-positive cells (r = −.78). The two ERα–PNN association measures were also strongly inversely related (r = −.95). Together, these exploratory findings indicate that the amount of maternal care, rather than its temporal unpredictability, covaries with the organization of PNNs around ERα-expressing cells following resource scarcity in the MPOA.

To determine whether the association between maternal care and ERα–PNN organization differed by housing condition, regression slopes were compared between CTRL and LBN dams. The slopes relating pup-directed behavior to the percentage of ERα-positive cells surrounded by PNNs did not differ between groups (F(1, 8) = 0.04, p = .84). Consistent with this result, the correlation coefficients were positive in both CTRL (r = .80) and LBN dams (r = .94). In contrast, the relationship between pup-directed behavior and the percentage of PNNs surrounding ERα-positive cells differed significantly between housing conditions (F(1, 8) = 8.60, Holm–Šídák-adjusted p = .037). This association was positive in CTRL dams (r = .50) but strongly negative in LBN dams (r = −.92). Thus, LBN altered how the relative distribution of PNNs around ERα-positive cells covaried with pup- directed maternal behavior, without altering the relationship between maternal care and the proportion of ERα-positive cells enwrapped by PNNs.

### LBN increases AR-positive cells while shifting remaining PNNs toward AR-expressing cells in the mPOA

Similar to what was observed for ERα, LBN increased the number of AR-positive cells (t(9) = 4.99, p < .001, η² = .73; Fig. 7A) and decreased PNN abundance (t(9) = 6.51, p < .001, η² = .82; Fig. 7B) compared with CTRL dams. As a result of these changes, the percentage of AR-positive cells surrounded by PNNs was significantly reduced in LBN dams (t(9) = 5.05, p < .001, η² = .74; Fig. 7C). In contrast to what was observed for ERα, the percentage of PNNs surrounding AR-positive cells was higher in LBN than in CTRL dams (t(9) = 7.90, p < .0001, η² = .87; Fig. 7D), suggesting that a greater proportion of the remaining PNNs in the mPOA of LBN dams was associated with AR-expressing cells.

**Figure 7.**
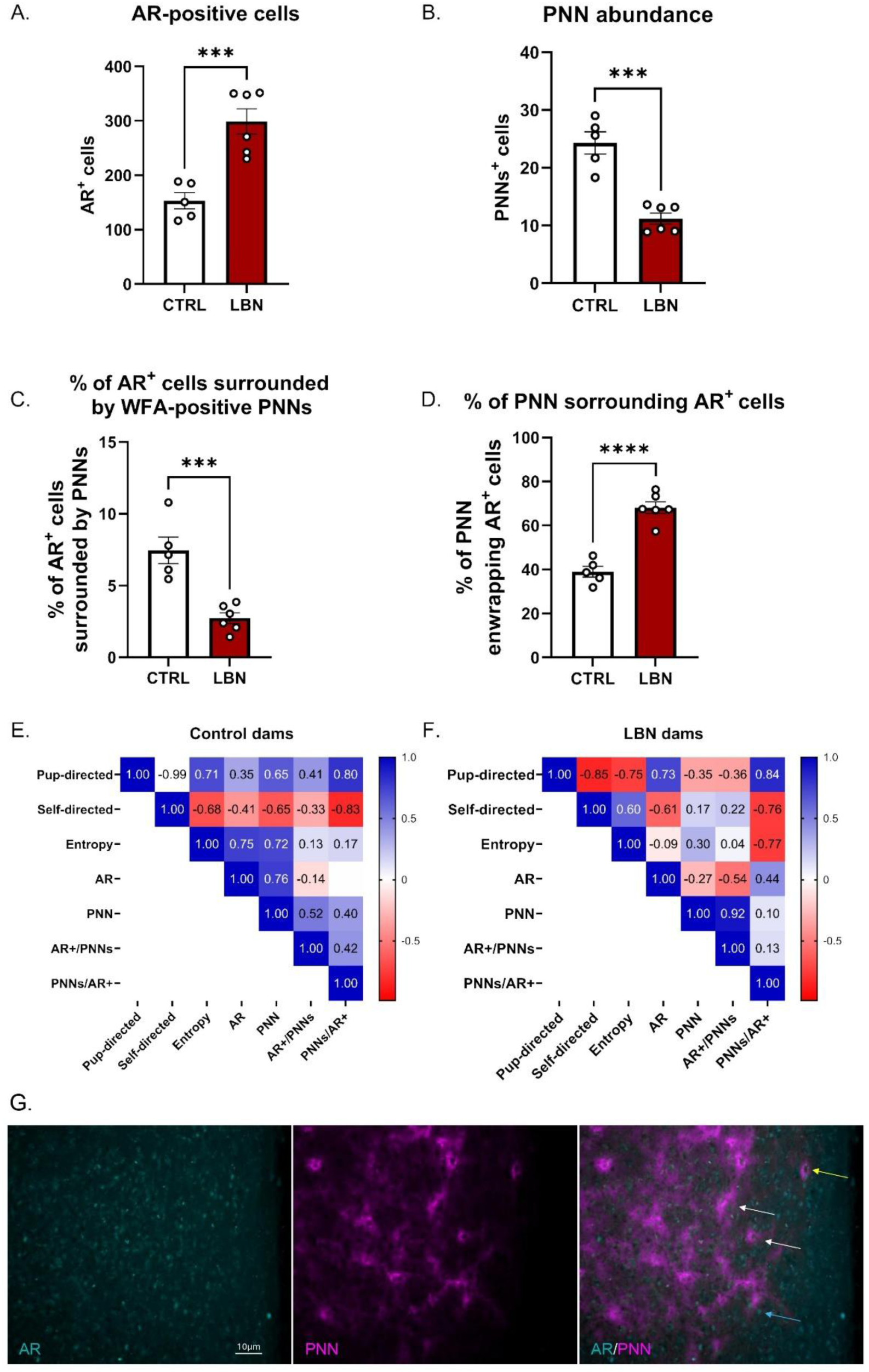
LBN increases AR-positive cells and shifts the distribution of remaining PNNs toward AR-expressing cells in the mPOA. (A) Number of androgen receptor-positive (AR+) cells in the medial preoptic area (mPOA). (B) Number of Wisteria floribunda agglutinin-labeled perineuronal nets (PNNs). (C) Percentage of AR+ cells surrounded by PNNs. (D) Percentage of PNNs surrounding AR+ cells. LBN increased the number of AR+ cells, reduced overall PNN abundance and the proportion of AR+ cells enwrapped by PNNs, and increased the proportion of the remaining PNNs associated with AR+ cells. (E, F) Exploratory Pearson correlation matrices showing associations among pup-directed behavior, self-directed behavior, behavioral entropy, AR+ cell number, PNN abundance, the percentage of AR+ cells surrounded by PNNs, and the percentage of PNNs surrounding AR+ cells in control (E) and LBN dams (F). (G) Representative fluorescence micrographs of AR immunoreactivity (cyan; left), WFA-labeled PNNs (magenta; center), and merged AR/PNN labeling (right) in the mPOA. In the merged image, white arrows indicate AR-positive cells enwrapped by WFA-positive PNNs, light- blue arrows indicate AR-positive cells without detectable PNN enwrapment, and yellow arrows indicate WFA-positive PNNs not enwrapping AR-positive cells. Values within the matrices represent Pearson correlation coefficients (r); blue indicates positive associations and red indicates negative associations. Formal comparisons of selected regression slopes did not reveal significant differences between housing conditions. Bars represent mean ± SEM, and individual points represent dams. n = 5 control and 6 LBN dams. Group comparisons in A–D were performed using independent-samples t-tests. ***p < .001; ****p < .0001. CTRL, control housing; LBN, limited bedding and nesting.

Exploratory correlation analyses indicated that associations among maternal behavior, AR expression, and PNN organization differed descriptively between CTRL and LBN dams (Fig. 7E, F). In CTRL dams, pup-directed behavior was positively associated with total PNN abundance (r = .65) and the percentage of PNNs surrounding AR-positive cells (r = .80), whereas self-directed behavior showed corresponding inverse associations (r = −.65 and r = −.83, respectively). In LBN dams, pup- directed behavior was positively associated with AR expression (r = .73) and remained positively associated with the percentage of PNNs surrounding AR-positive cells (r = .84). However, its associations with total PNN abundance (r = −.35) and the percentage of AR-positive cells surrounded by PNNs (r = −.36) were negative.

Behavioral entropy was positively associated with AR expression (r = .75) and PNN abundance (r = 0.72) in CTRL dams. These associations were weaker in LBN dams (r = −.09 and r = .30, respectively), in which entropy was instead negatively associated with the percentage of PNNs surrounding AR-positive cells (r = −.77). AR expression and PNN abundance were also positively associated in CTRL dams (r = .76) but negatively associated in LBN dams (r = −.27). However, direct comparisons of the selected regression slopes found no significant differences between housing conditions (all p > .05). Thus, although the correlation matrices showed descriptive differences, these analyses did not provide evidence that LBN altered the relationships among maternal behavior, AR expression, and PNN organization.

### Corticosterone increases in pup plasma, but only at the end of the procedure and in females

Plasma corticosterone concentrations were measured in pups from independent litters at PND6 and PND10. At PND6, two-way ANOVA revealed no significant effect of housing condition (F(1, 31) = 1.65, p = .21), pup sex (F(1, 31) = 0.02, p = .90), or housing condition × sex interaction (F(1, 31) <1; Fig. 8A). At PND10, two-way ANOVA revealed a significant main effect of housing condition (F(1, 30) = 12.65, p = .0013, η² = 0.25), no main effect of pup sex (F(1, 30) = 0.009, p = .92, η² < .001), and a significant housing condition × sex interaction (F(1, 30) = 7.32, p = .011, η² = .15; Fig. 8B). Šídák- adjusted comparisons showed that female pups from LBN litters had higher plasma corticosterone concentrations than CTRL female pups (p < .001), whereas male pups did not differ between housing conditions (p > .05).

**Figure 8.**
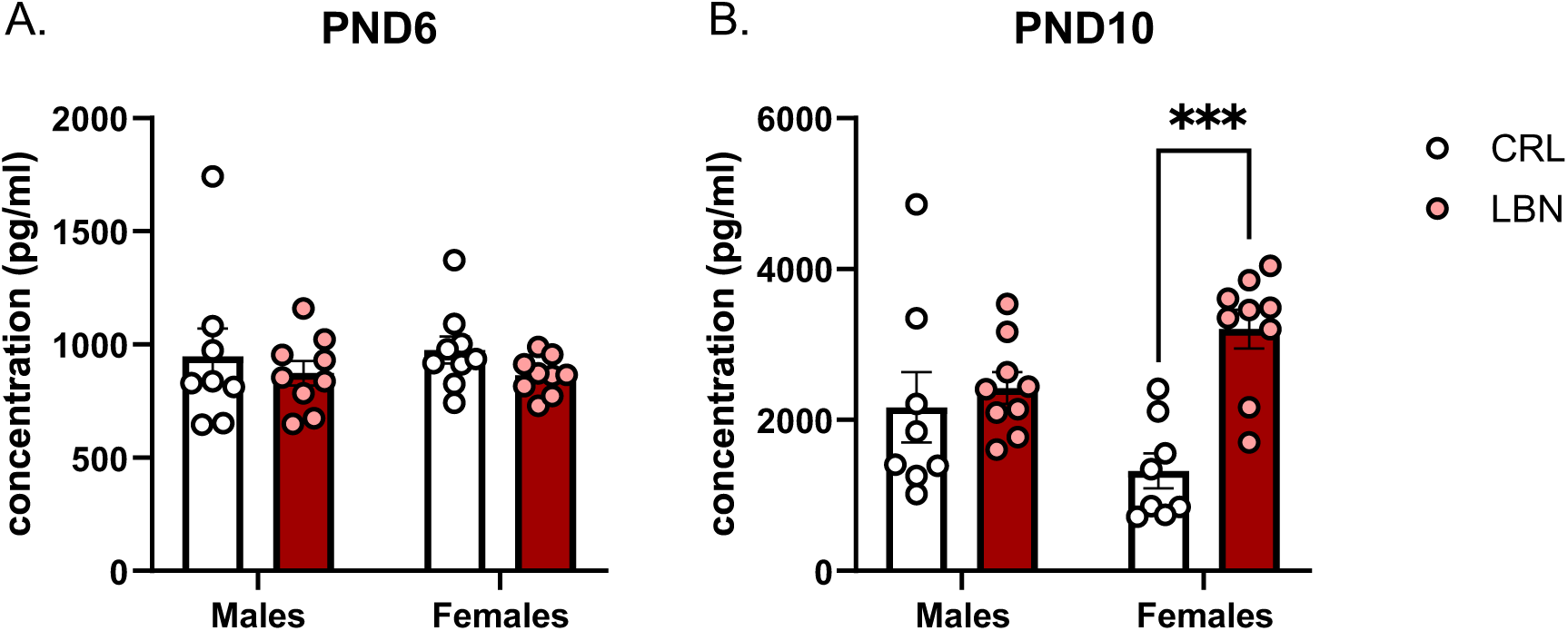
LBN increases plasma corticosterone selectively in female pups at PND10. Plasma corticosterone concentrations were measured by ELISA in male and female pups from CTRL and LBN litters at (A) PND6 and (B) PND10. No significant effects were detected at PND6. At PND10, Šídák- adjusted comparisons showed higher corticosterone concentrations in LBN female pups than in CTRL female pups, whereas males did not differ between housing conditions. Bars represent mean ± SEM, and symbols represent individual pups, each derived from a different litter. A total of 35 pups/independent litters were analyzed at PND6 and 34 at PND10. Sample sizes were PND6: CTRL males, (n = 8); CTRL females, (n = 9); LBN males, (n = 9); and LBN females, (n = 9); and PND10: CTRL males, (n = 8); CTRL females, (n = 8); LBN males, (n = 9); and LBN females, (n = 9). Each pup originated from a different litter and represents an independent biological replicate. **p < .001.

## Discussion

The present study demonstrates that postpartum resource scarcity reorganizes maternal behavior across multiple levels, from the temporal architecture of caregiving to the allocation of responses among offspring and the structural organization of steroid-sensitive circuits within the mPOA. Importantly, these effects were not consistent with a simple reduction in maternal care. Instead, LBN dams spent more time engaged in pup-directed behaviors than control dams and less time in self-directed activities, indicating sustained or even enhanced engagement with offspring. At the same time, caregiving became less predictable and more fragmented, retrieval was completed less efficiently, and behavioral responses became selectively biased toward male pups. These behavioral changes were accompanied by increased numbers of ERα- and AR-immunoreactive cells, reduced PNN abundance, and receptor-specific changes in PNN association within the mPOA, suggesting that resource scarcity selectively reorganizes extracellular-matrix constraints on experience-dependent plasticity across steroid-sensitive maternal circuits. Female, but not male, pups also exhibited increased plasma corticosterone at PND10. Together, these findings suggest that scarcity alters how maternal effort is allocated rather than whether maternal care is expressed, and suggests this may affect stress levels in pups.

The coexistence of increased pup-directed care and increased behavioral entropy reinforces the idea that the quantity of care is separable from its organization. Much of the literature frames maternal care along a quantitative dimension, with adversity viewed primarily as a factor that increases or decreases the total amount of care provided (Baram et al., 2012). Previous studies differ in how scarcity affects caregiving quantity. In Sprague–Dawley rats, Ivy et al. (Ivy et al., 2008) reported reduced licking/grooming and more frequent departures from the nest, whereas Gallo et al. (Gallo et al., 2019) found increased nest attendance alongside more frequent nest entries and exits in C57BL/6 mice. More closely resembling the present findings, studies have reported increased pup- directed behaviors, reduces self-directed ones and more behavioral transitions (Pardoet al., 2024; Shupe C Clinton, 2021). Although transition frequency and entropy quantify different features of behavioral organization, these observations, together with the increased maternal behavioral entropy reported by Demaestri et al. (Demaestri et al., 2022), argue against treating temporal disruption as synonymous with reduced caregiving. Differences in protocols, species, strain, genetic background, observation period, and behavioral definitions may contribute to the contrasting effects on care quantity (Demaestri et al., 2022; Gallo et al., 2019; Ivy et al., 2008; Pardo et al., 2024). In the present study, increased pup-directed behavior and reduced self-directed activity are consistent with a redistribution of the observed maternal time budget toward offspring. This pattern could reflect compensatory or hypervigilant responding to an unstable environment, as proposed by Gallo et al. (Gallo et al., 2019). However, increased contact should not be equated with successful compensation. Indeed, under a later limited-bedding protocol spanning PND8–12, Song et al. (Song et al., 2026) found that sustained nest presence coexisted with increased rough handling and less efficient pup transport. Thus, substantial maternal engagement can coexist with compromised aspects of caregiving. Elevated entropy establishes lower predictability of behavioral sequences, but does not by itself identify diminished motivation, inefficient decision-making, or an adaptive response to competing demands. Our findings therefore support a reorganization of caregiving under constraint, while leaving unresolved whether increased pup-directed time compensates for, or accompanies, impaired temporal coordination.

Beyond altering the amount and temporal organization of maternal care, resource scarcity also shifted which offspring dams prioritized. Relative to CTRL, LBN dams were more likely to retrieve a male first and allocated more investigation and goal-arm time toward males when male and female pups were presented together. These differences occurred without corresponding changes in general Y-maze exploration, arguing against a generalized change in activity and instead suggesting altered prioritization of offspring-related information. This interpretation extends evidence that pups are distinguishable, multimodal stimuli: male and female pups can differ in vocal output and retrieval order, as well as in urinary odors that dams discriminate (Bowers et al., 2013; McRae et al., 2023; Moore, 1985). Moreover, LBN can alter pup temperature, movement, and vocal behavior, indicating that adversity changes the information pups provide to their caregivers (Lapp et al., 2024). Scarcity could therefore influence maternal choices by changing pup-derived information, the maternal weighting of that information, or both.

The adaptive significance of this male bias remains unresolved. Comparative research demonstrates that environmental conditions can alter which offspring signals guide parental provisioning: in birds, poor or unpredictable environments are associated with reduced responsiveness to begging and greater reliance on structural cues of offspring quality (Caro et al., 2016), while theoretical models similarly predict that provisioning decisions should depend on environmental quality (J. N. Davis et al., 1999) . Nevertheless, adversity does not uniformly promote male-directed care. Power and Moore (Power C Moore, 1986) found that the preferential licking normally received by male rat pups was absent among prenatally stressed pups; urine from stressed males also elicited less maternal investigation, suggesting that altered pup chemosignals contributed to this effect. Although prenatal stress and LBN differ in timing and licking and retrieval represent distinct components of care, this contrast demonstrates that sex-biased maternal responses depend on the caregiving context. Moreover, classical sex-allocation theory does not straightforwardly predict the male bias observed here: predictions concerning offspring sex ratio differ from those concerning investment within an existing mixed-sex litter, and the latter require additional assumptions about sex-specific fitness returns (Trivers C Willard, 1973; Veller et al., 2016). Because our assays measured brief retrieval and investigation choices rather than cumulative care or reproductive outcomes, the strongest conclusion is that LBN shifted maternal prioritization toward male-associated, multimodal information under these testing conditions and not that males were intrinsically valued more highly or that this shift was adaptive.

The female-biased increase in pup corticosterone at PND10 identifies a sex-dependent physiological outcome of LBN. One possibility is that the male-biased maternal responses observed here reflect a redistribution of care within the litter, leaving female pups exposed to less or differently organized maternal input. This interpretation is appealing because the endocrine difference occurred in the sex that was relatively deprioritized across the behavioral tasks, but it remains speculative. Future studies during home-cage care and linking pup-specific caregiving histories to corticosterone levels are needed. The timing of this effect is nevertheless notable because PND10 falls within the conventional stress-hyporesponsive period in rats, approximately PND4–14, during which basal corticosterone is generally low and responses to mild stressors are attenuated. Maternal inputs, including feeding and tactile stimulation, contribute to maintaining this hyporesponsive state (Okimoto et al., 2002; Suchecki et al., 1993) . The elevation in LBN females could therefore reflect a sex-specific alteration in HPA-axis regulation or maternal buffering of males during this developmental window. However, the stress-hyporesponsive period is relative rather than absolute, and its disruption cannot be established from a single corticosterone measurement. LBN-related corticosterone phenotypes vary with strain, protocol, developmental stage, and sampling conditions; reduced rather than increased corticosterone has also been reported at PND10 under other limited-nesting procedures (Moussaoui et al., 2016; Walker et al., 2017). Nevertheless, the female-selective elevation observed here could represent an early endocrine correlate of later sex- divergent outcomes, including reduced vascular coverage in the only in medial prefrontal cortex of females (Andrews et al., 2026) and male-specific protection from some addiction-related phenotypes previously reported in this implementation of LBN (Sanchez et al., 2021). If persistent or recurrent, altered corticosterone exposure could influence the development of stress- and reward- related circuits. Future work establishing the duration and functional consequences of this effect are needed.

The anatomical findings indicate that this behavioral reorganization was accompanied by remodeling of a steroid-sensitive maternal circuit. The mPOA integrates pup-derived sensory information with endocrine and physiological state and is central to parental behavior (Jamieson C Kohl, 2026; Numan C Insel, 2006). ERα signaling in this region is required for normal pup retrieval and covaries with maternal care, whereas AR-positive populations in maternal females remain less understood (Champagne et al., 2003; Tsuneoka et al., 2017; Wei et al., 2018). LBN increased ERα- and AR-immunoreactive cell counts, which may indicate altered receptor expression or detectability rather than generation of new cells. LBN also reduced PNN abundance and the proportion of both receptor-positive populations enwrapped by PNNs. Previous work demonstrated that mPOA PNNs remodel across pregnancy and lactation and can surround ERα- and progesterone receptor-positive cells (Uriarte et al., 2020). The present study extends these findings by showing that the postpartum environment further alters PNN abundance and cellular distribution and, to our knowledge, provides the first characterization of PNN association with AR-positive cells in the maternal mPOA. The pattern was not identical across receptor-defined analyses: the proportion of WFA-positive PNNs surrounding AR-positive cells increased, whereas the corresponding ERα measure was unchanged. This pattern is consistent with a relative concentration of the remaining detectable nets around AR- positive cells. Experimental degradation or attenuation of PNNs can alter experience-dependent synaptic plasticity and behavior in adult circuits, including recognition memory and fear-related learning (Gogolla et al., 2009; Hylin et al., 2013; Romberg et al., 2013). Together, our findings suggest that postpartum resource scarcity reorganizes the extracellular-matrix environment of steroid- sensitive mPOA populations, potentially altering their capacity for experience-dependent remodeling and adaptation to offspring-related demands.

One possible molecular route is steroid-sensitive regulation of extracellular-matrix turnover. Estradiol can alter PNN labeling and the expression of genes encoding matrix metalloproteinases (MMPs), a disintegrin and metalloproteinase with thrombospondin motifs (ADAMTS) enzymes, and their endogenous inhibitors in the female brain; it can also rapidly increase hippocampal MMP-9 activity (Gross et al., 2022; Nguyen et al., 2025). These findings make MMP-9 a plausible candidate, but they do not establish an ERα–MMP-9 pathway in the maternal mPOA. Likewise, whether neural AR signaling regulates PNN turnover through endogenous metalloproteinase inhibitors or specific ADAMTS enzymes remains unknown. A parsimonious model is that scarcity shifts the maternal system toward high pup contact but less stable behavioral organization while changing the weighting of offspring-related information, accompanied by reorganization of the steroid-sensitive extracellular matrix in the mPOA. This model is compatible with evidence that ERα-expressing mPOA neurons promote pup approach and retrieval and that pregnancy hormones remodel galanin- expressing mPOA neurons to enhance responses to pups (Ammari et al., 2023; Fang et al., 2018). Nevertheless, the anatomical and behavioral effects could be parallel consequences of altered endocrine, metabolic, or sensory conditions. Establishing causality will require manipulating ERα- or AR-defined mPOA populations and PNN integrity while assaying matrix-remodeling pathways and measuring both behavioral sequence structure and offspring-sex prioritization.

In summary, postpartum resource scarcity did not simply suppress maternal care. It increased the time dams devoted to pups while making caregiving less predictable, delaying completion of retrieval, and shifting maternal responses toward male offspring. These changes were accompanied by receptor-specific reorganization of PNN architecture in the mPOA in dams and a female-specific corticosterone elevation at PND10 in pups. The translational importance of caregiving predictability is underscored by longitudinal human studies linking greater unpredictability of maternal sensory signals during infancy to poorer cognitive and episodic-memory outcomes, altered corticolimbic maturation, and blunted cortisol reactivity (Davis et al., 2017; Granger et al., 2021; Noroña-Zhou et al., 2020). To our knowledge, the present study provides the first evidence that LBN alters PNN architecture in the maternal mPOA and reorganizes its association with ERα- and AR-positive populations, identifying steroid-sensitive extracellular-matrix remodeling as a candidate neural substrate accompanying the reorganization of care under adversity. Increased pup- directed behavior may represent a compensatory response that helps sustain offspring care under low-resource conditions; however, its fragmentation and preferential allocation toward males suggest that its potential stress-buffering effects may not be distributed equally across the litter. Thus, maternal stress may shape offspring vulnerability not only by diminishing care, but by reorganizing when and toward whom care is allocated, which may potentially translate environmental scarcity into sex-different developmental outcomes.

## Acknowledgment

This work was supported by the National Institute on Drug Abuse (R01 DA049837, R01 DA056534, and R21 DA062844 to D.B.), the Wilczynski–Georgia Research Alliance Future Faculty Fellowship Program (to D.A.), and a BBRF Young Investigator Grant (Award No. 34513 to D.A.). We thank Sydney Ku for assistance with formatting the transition probability matrices and Ana Isabel Silva for assistance with confocal imaging. Figures were created using BioRender.com.

